# Mechanistic Insights into the Protective Roles of Polyphosphate Against Amyloid Cytotoxicity

**DOI:** 10.1101/704882

**Authors:** Justine Lempart, Eric Tse, James A. Lauer, Magdalena I Ivanova, Alexandra Sutter, Nicholas Yoo, Philipp Huettemann, Daniel Southworth, Ursula Jakob

## Abstract

The universally abundant polyphosphate (polyP) accelerates fibril formation of disease-related amyloids and protects against amyloid cytotoxicity. To gain insights into the mechanism(s) by which polyP exerts these effects, we focused on α-synuclein, a well-studied amyloid protein, which constitutes the major component of Lewy bodies found in Parkinson’s Disease. Here we demonstrate that polyP is unable to accelerate the rate-limiting step of α-synuclein fibril formation but effectively nucleates fibril assembly once α-synuclein oligomers are formed. Binding of polyP to α-synuclein either during fibril formation or upon fibril maturation substantially alters fibril morphology, and effectively reduces the ability of α-synuclein fibrils to interact with cell membranes. The effect of polyP appears to be α-synuclein fibril specific, and successfully prevents the uptake of fibrils into neuronal cells. These results suggest that altering the polyP levels in the extracellular space might be a potential therapeutic strategy to prevent the spreading of the disease.

## INTRODUCTION

Parkinson’s disease (PD), the second most common neurodegenerative disorder known ^1^, is characterized by a loss of dopaminergic neurons in the *substantia nigra* ^2, 3^. A hallmark of the disease is the appearance of intracellular protein inclusions (i.e., Lewy bodies), which consist primarily of insoluble fibrils of α-synuclein, a 140-aa protein involved in presynaptic vesicle formation ^4, 5^. While it is now well established that deposition of α-synuclein fibrils associates with the disease and that cell death can be elicited simply by incubating neuronal cells with α-synuclein fibrils ^6^, many open questions remain concerning the mechanism of toxicity, the structural features of the toxic α-synuclein species and the way(s) by which α-synuclein toxicity propagates in the brain.

In solution, α-synuclein is a soluble monomer with extensive regions of intrinsic disorder ^7^. *In vitro* studies demonstrated that upon prolonged incubation, α-synuclein monomers undergo conformational rearrangements, which lead to the formation of aggregation sensitive oligomers ^8^. These nuclei are capable of sequestering other α-synuclein monomers and will grow into proto-fibrils and eventually into insoluble, protease-resistant fibrils ^9, 10^. *In vitro*, the rate-limiting step in fibril-formation appears to be the formation of the initial nuclei, and fibril formation has been shown to be accelerated by the addition of negatively charged polymers, including glucosaminoglycans (i.e., heparin) ^11^, RNA ^12^ or phospholipids ^13^. The precise roles that these additives play in *in vivo* fibril formation remain to be determined.

Recent studies provided supporting evidence that amyloid toxicity is not caused by the fibrils *per se* but by oligomeric species that transiently accumulate on the pathway to fibril formation ^6, 14^. These oligomers, which have been shown to affect mitochondrial function ^15^, membrane permeability ^16, 17^ and/or the cytoskeleton ^18^, are thought to be responsible for the observed neuroinflammation ^19^ and cell death ^6^. Moreover, amyloid oligomers seem to be the primary species that spread among cells ^20, 21^, and to be responsible for the prion-like propagation of PD pathology ^22, 23^. Cell-to-cell transmission appears to involve the active secretion of α-synuclein oligomers into the extracellular space followed by the uptake of the amyloids into neighboring recipient cells via micropinocytosis and glucosaminoglycan (GAG) receptors ^24–26^. Experiments conducted in cell culture confirmed that α-synuclein oligomers can readily spread between neurons and glial cells ^27, 28^, and, once taken up by recipient cells, sequester monomeric α-synuclein into insoluble foci ^25, 29^.

Recent work from our lab demonstrated that polyphosphate (polyP), a highly conserved and universally present polyanion, significantly decreases the cytotoxicity of amyloidogenic proteins ^30^. These results were corroborated in studies with amyloid β_25-35_, which showed that pre-incubation of PC12 cells or primary cortical neurons with polyP protects against the neurotoxic effects of the peptide ^31^. *In vitro* studies revealed that polyP substantially accelerates α-synuclein fibril formation in a chain-length dependent manner, causing the formation of both shedding-resistant and seeding-deficient polyP-associated fibrils ^30^. Localization studies revealed that polyP, like α-synuclein, is both secreted and taken up by neuronal cells and hence localizes both inside and outside of cells ^32^. These results raised intriguing questions as to what α-synuclein species interact with polyP, how premature fibril formation might be avoided, and, most importantly, by what mechanism polyP is able to protect neuronal cells against α-synuclein toxicity.

Here we show that polyP does not interact with monomeric α-synuclein but effectively nucleates α-synuclein fibril formation once prefibrillar species are present. PolyP causes pronounced morphological changes in both *de novo* forming fibrils as well as upon its addition to mature α-synuclein fibrils, demonstrating that α-synuclein fibrils are inherently dynamic and amendable to polyP-mediated structural changes. Importantly, presence of polyP strongly interferes with the interaction of α-synuclein fibrils with cell membranes, and prevents the uptake of α-synuclein fibrils into differentiated neuroblastoma cells. These results explain the cytoprotective effect of polyP and suggest that extracellular polyP might be able to influence the spreading of this disease.

## RESULTS

### PolyP accelerates fibril formation by nucleating α-synuclein oligomers

Amyloid fibril formation is most commonly monitored by measuring the fluorescence of thioflavin T (ThT), a small molecular dye that becomes highly emissive when intercalated into the β-sheets of amyloidogenic oligomers and fibrils ^33^. ThT kinetics of amyloid fibril formation can be divided into three distinct phases (Fig. 1a); the nucleation (i.e. lag) phase, in which soluble monomers undergo structural changes and nucleate; the elongation (i.e., growth) phase, during which ThT-positive oligomers and proto-fibrils form; and the equilibration (i.e., plateau) phase, in which mature fibrils undergo cycles of shedding and seeding ^34^. Consistent with our previous *in vitro* studies ^30^, we found that polyP substantially accelerates the fibril-forming process, both by shortening the lag phase and by increasing the rate of fibril growth (Fig. 1b, c). To determine when during the polymerization process polyP acts on amyloidogenic proteins, we conducted fluorescence polarization (FP) measurements, which record the tumbling rate of fluorescent molecules as read-out for real-time binding events ^35^. We labeled polyP_300_-chains (MW: ~30 kDa) with AlexaFluor 647 (polyP_300-AF647_) and conducted FP-measurements in the presence of freshly prepared α-synuclein (MW: 14 kDa) for 40 hours (Fig. 1b, middle panel). Unexpectedly, we did not observe any significant increase in the FP-signal over the first ~2.5 h of incubation, suggesting that polyP does not interact with monomeric α-synuclein. After this lag-phase, however, the FP-signal rapidly increased and reached a plateau after about 12 h of incubation. By normalizing the ThT-aggregation and FP-binding results, we observed that the increase in FP-signal (T_1/2_ = 5.1 ± 1.4h) may slightly precede the formation of ThT-positive amyloid intermediates (T_1/2_ = 8.5 ± 1.6h) (Fig. 1b, insert in right panel). These results suggested that polyP does not interact with α-synuclein species that occur early in the fibril-forming process (i.e., monomers) but instead binds α-synuclein species shortly before or concomitant with their ability to intercalate ThT. Time-delayed polyP addition experiments confirmed these results and demonstrated that polyP acts on nucleation-competent oligomers and/or proto-fibrils. For these studies, we used experimental conditions under which α-synuclein fibril formation proceeds with a lag phase of ~ 6h in the absence and ~ 1.5 h in the presence of polyP (Fig. 1c, compare open and cyan circles). When we added polyP two hours after the start of the incubation, the lag-phase was reduced from the remaining 4h in the absence of polyP to less than 30 min (Fig. 1c, green squares). Addition of polyP after 5h caused an immediate increase in ThT signal (Fig. 1c, blue diamonds) while addition of polyP mid-way through the elongation phase of α-synuclein fibril formation triggered maximal ThT binding within less than 10 min (Fig. 1c, red triangles). These results strongly suggested that polyP binds to a range of presumably non-monomeric α-synuclein species and supports their association into insoluble fibrils.

**Figure 1:**
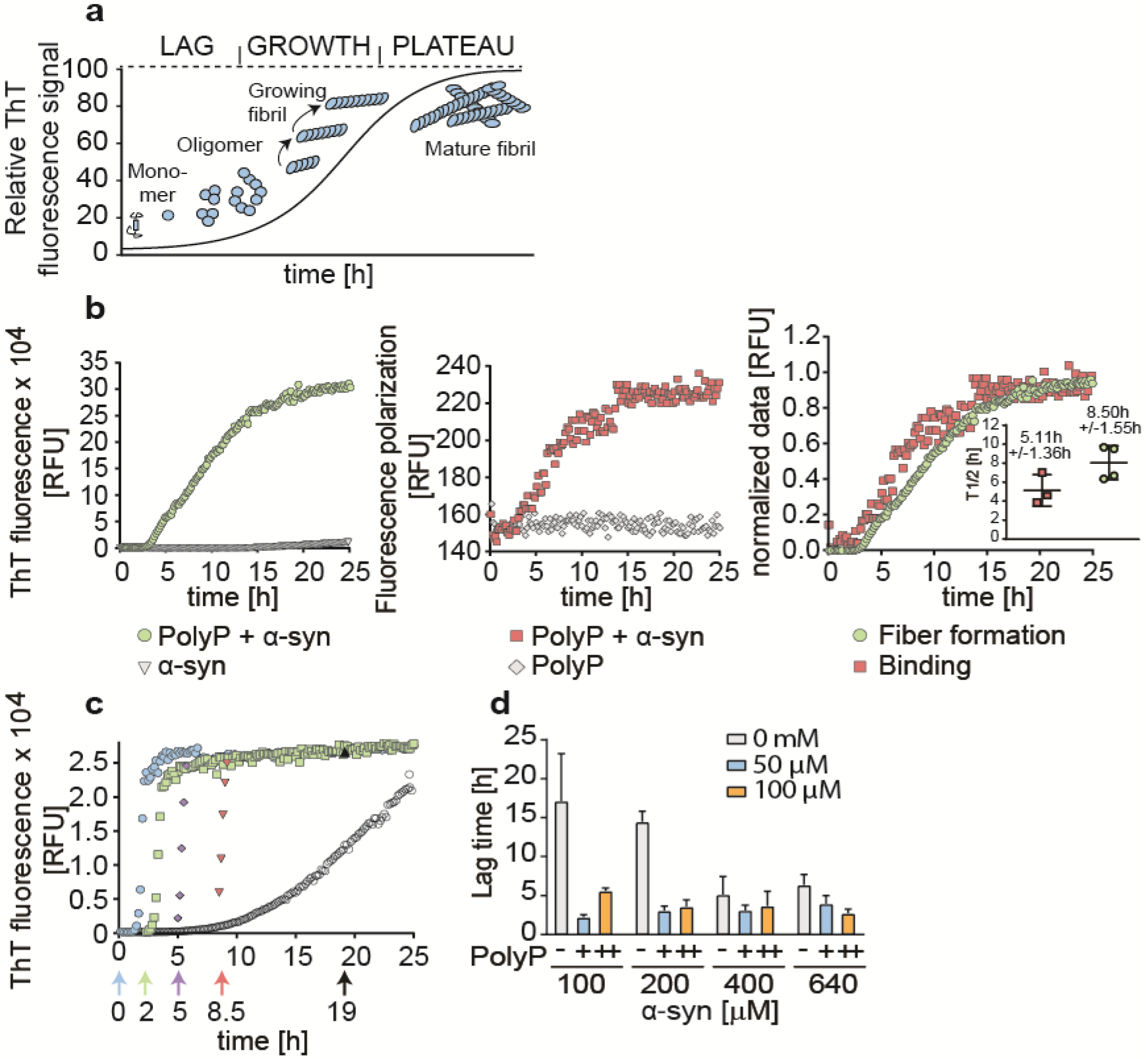
Influence of polyP on α-synuclein fibril formation *in vitro*. **a.** Model of the amyloid fibril forming process using ThT-fluorescence. **b.** 100 μM freshly prepared α-synuclein was incubated in the absence or presence of 500 μM fluorescently labeled polyP_300-AF647_ (in P_i_ units) at 37°C under constant stirring. ThT fluorescence was used to monitor fibril formation (left panel) and fluorescence polarization (FP) experiments were conducted to measure binding of polyP_300-AF647_ to α-synuclein (middle panel). Overlay of normalized ThT and FP-curves (right panel). Insert: Half time (T_1/2_) of fibril formation and fluorescence polarization measurements. Data are the mean of three independent experiments +/- SD. **c**. Addition of 500 μM polyP (in P_i_ units) before (cyan circles) or at defined time-points during the fibril-forming process of 300 μM α-synuclein. ThT fluorescence was monitored. All experiments were conducted at least three times. Representative kinetic traces are shown. **d**. Influence of different polyP_300_ and α-synuclein concentrations on the lag phase of fibril formation. ThT fluorescence was monitored and the lag phase was determined. The mean of four experiments +/- SD is shown.

### PolyP does not affect rate-limiting step of fibril formation

Our finding that polyP does not detectably interact with α-synuclein monomers but readily stimulates fibril formation once ThT-positive oligomers have formed, suggested that polyP does not affect the rate-limiting step of α-synuclein fibril formation. To test this idea, we combined increasing α-synuclein concentrations with increasing polyP concentrations, and measured the respective lag phase of fibril formation using ThT fluorescence (Fig. 1d). As expected, increasing the α-synuclein concentration from 100 μM to 400 μM in the absence of polyP reduced the lag phase from about 16 h to less than 7 h. Higher concentrations of α-synuclein (i.e., 640 μM) did not significantly shorten the lag phase any further. The presence of physiological relevant concentrations of polyP_300_ (50 μM in P_i_-units) ^36, 37^ reduced this lag time to 2-3 h. Noteworthy, this reduction in lag time appeared to be independent of the α-synuclein concentration used (Fig. 1d). Moreover, doubling the polyP concentration also failed to further reduce the lag phase. These results agreed with previous results showing that α-synuclein undergoes conformational changes and/or oligomerization processes that are rate-limiting ^9, 38^, and suggested that this step cannot be accelerated by the presence of polyP. We concluded from these results that simple co-existence of polyP and α-synuclein in the same (extra)cellular compartment will unlikely be sufficient to trigger *de novo* fibril formation.

### PolyP alters morphology of pre-formed α-syn fibrils

FP-binding studies using pre-formed α-synuclein fibrils revealed that polyP not only interacts with ThT-positive oligomers during *de novo* fibril formation but also binds to mature fibrils (Fig. 2a). Since α-synuclein fibrils that are formed in the presence of polyP (i.e., α-syn^polyP^) have significantly altered morphology compared to fibrils formed in the absence of nucleators (i.e., α-syn^alone^) ^30^, we wondered whether polyP binding would also affect the morphology of mature fibrils. This would possibly explain why the addition of polyP to preformed fibrils was as cytoprotective as its addition during fibril formation ^30^. We therefore generated α-synuclein fibrils, washed and purified them to remove any small oligomers and protofibrils, and either left them untreated (α-syn^alone^) or incubated them with polyP_300_ (α-syn^alone→polyP^). Immediately before as well as 20 min after the addition of polyP to α-syn^alone^ fibrils, we fixed aliquots of the samples on grids, and prepared them for transmission electron microscopy (TEM). As a control, we also tested α-syn fibrils formed in the presence of polyP (α-syn^polyp^). As shown in Fig. 2b, the morphology of α-syn^alone→polyP^ fibrils was nearly indistinguishable from the morphology of α-syn^polyP^ fibrils. Instead of two protofilaments, which typically form a twisted structure, α-syn^alone→polyP^ and α-syn^polyP^ fibrils were significantly thinner, suggesting that polyP caused their dissociation into single protofilaments. X-ray fibril diffraction measurements agreed with the finding that incubation of preformed fibrils with polyP alters their conformations, and showed particularly striking differences in the equatorial plots of the radial intensities (i.e., X-axis), which arise from the packing of adjacent β-sheets in the amyloid fibril. In contrast, no differences were observed on the meridian (Y-axis), which reflects the strand-to-strand packing, and produced a sharp reflection at 4.7 Å spacing for both fibril species (Fig 2c, d). These results suggested a pronounced effect of polyP on the packing of the β-sheets within the protofilament (Fig. 2e).

**Figure 2:**
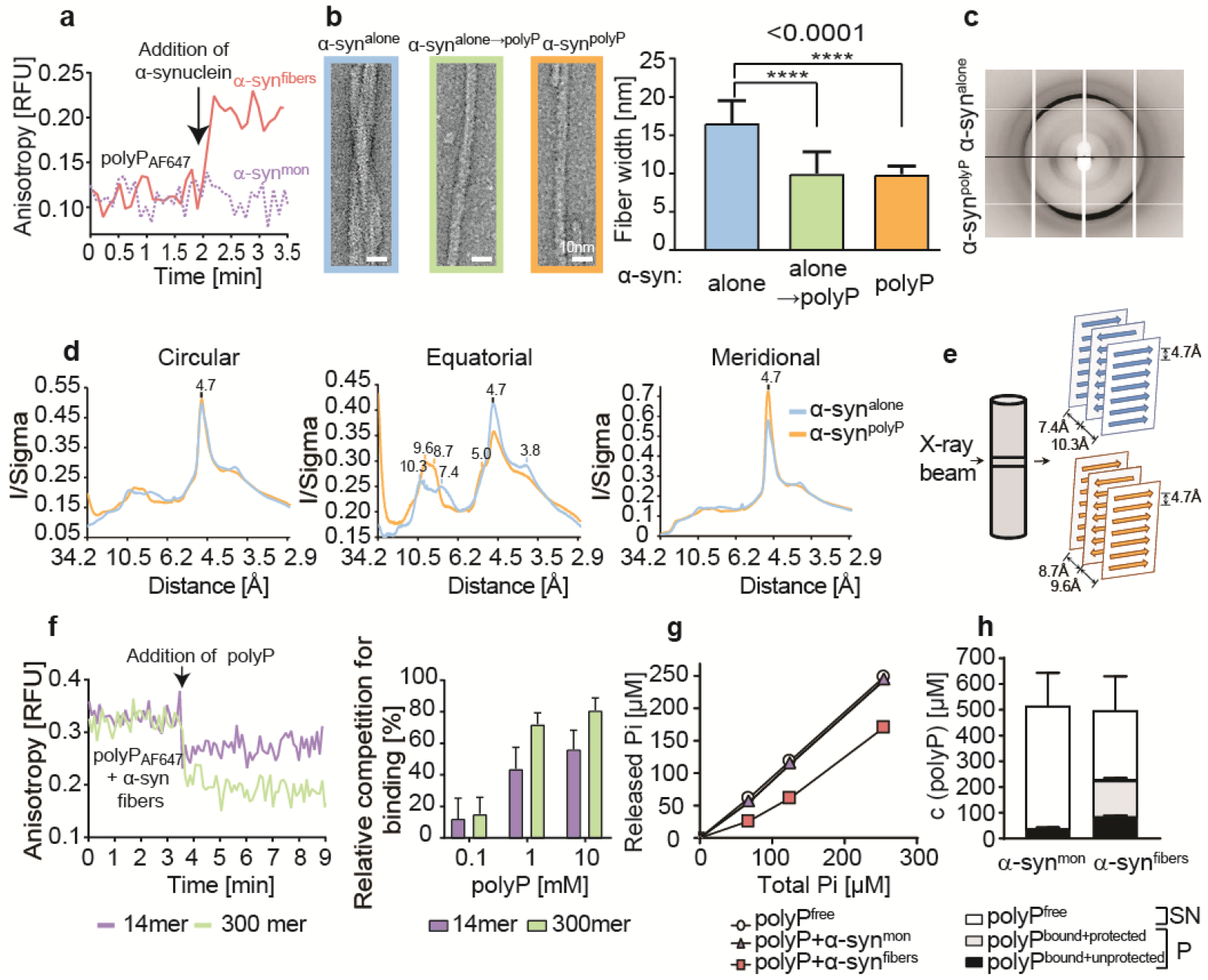
Effects of polyP on α-synuclein fibril morphology. **a.** Fluorescence polarization of 50 μM polyP_300-AF647_ upon addition of 30 μM α-syn^mon^ or α-syn^fibrils^. The arrow indicates the time point of protein addition. **b.** Transmission electron microscopy of α-synuclein fibrils (300 μM) formed in the absence of polyP and left untreated (α-syn^alone^) or incubated with 7.5 mM polyP_300_ for 20 min at RT (α-syn^alone→polyP^). Alpha-synuclein fibrils formed in the presence of 7.5 mM polyP_300_ were used as control (α-syn^polyP^). Quantitative analysis of fibril width was based on 10 individual micrographs and about 45 individual α-syn filaments. Statistical analysis was prepared with ONE-way Annova (****; p-value <0.0001)**. c. and d.** X-ray fiber diffraction of α-synuclein formed in the absence or presence of polyP_300._ The oriented samples produced cross-β diffraction patterns that contained a sharp reflection at 4.7 Å spacing at the meridian (Y-axis) and a broad reflection ~9Å spacing at the equator (X-axis) (**c**). The intensities were radially averaged over a full circle (360°, left panel), an equatorial arc (±30° around X-axis, middle panel), and meridional arc (±30° around Y-axis, right panel) **(d**). **e.** Cartoon representation of the possible ways β-sheets and strands assemble in the fibril. **f.** Left panel: Fluorescence polarization of 30 μM of pre-formed α-synuclein - polyP_300-AF647_ fibrils before and after the addition of 1 mM unlabeled polyP_300_ or polyP_14_. The arrow indicated the time point of polyP addition. Right panel: Varying concentrations of polyP_14_ or polyP_300_ in the competition experiment. The percent competition was calculated from the relative signal change upon polyP addition, setting the polyP_300-AF647_ fibril signal to 0 % competition and the polyP_300-AF647_ alone signal as 100 % competition. The mean of three experiments +/- SD is shown. **g.** 40 μM α-synuclein monomers (triangles) or preformed fibrils (squares) were incubated with increasing concentrations of polyP_300_ for 10 min. Samples were treated with ScPPX and assayed for released P_i_. A standard curve of polyP_300_ in the absence of α-synuclein (circles) was used as control. **h.** 40 μM α-synuclein monomers or preformed fibrils were incubated with 500 μM polyP_300_ for 10 min. Samples were separated into supernatant (SN) and pellet (P), treated with ScPPX and subsequently assayed for P_i_. PolyP_bound-unprotected_ represents the amount of P_i_ that was released upon ScPPX treatment in the pellet fraction. The amount of polyP not released by ScPPX was considered to be protected by the fibrils against hydrolysis (polyP_bound + protected_).

To further investigate the dynamics of polyP-fibril interactions, we conducted FP-competition experiments with pre-formed α-syn-polyP_300-AF647_ fibrils (Fig. 2f). As expected, we observed a high initial FP-signal, consistent with the slow tumbling rate of polyP-fibril complexes. Upon addition of un-labeled polyP_300_, however, the FP-signal rapidly decreased, indicating that the unlabeled polyp-chains replaced the labeled polyP in the fibrils. Addition of the much shorter polyP_14_ chain also reduced the FP-signal but to a lesser extent, suggesting that shorter chains have lower binding affinities than longer chains (Fig. 2f). These results indicated that the polyP-fibril interactions are highly dynamic in nature, and implied that fibrils, even when formed in the absence of polyP, can rapidly adopt a novel conformation when exposed to polyP.

### PolyP-fibril complexes are polyphosphatase resistant

Unbound polyP is very rapidly degraded by exopolyphosphatases, such as yeast PPX, which hydrolyzes the phosphoanhydride bonds with a turnover rate of 500 μmol/mg/min at 37°C ^39^. To test whether degradation of polyP reverses the morphological changes that we observed in fibrils bound to polyP, we incubated α-syn^polyP^ fibrils with yeast PPX. Surprisingly, however, we did not observe any morphological changes in the α-synuclein fibrils by TEM (data not shown). These results suggested either that the fibrils maintain their altered conformation even upon hydrolysis of polyP or that polyP, once in complex with fibrils, resists PPX-mediated hydrolysis. To investigate whether PPX is able to degrade fibril-associated polyP, we incubated 40 μM α-synuclein monomers or preformed α-synuclein fibrils with increasing concentrations of polyP, added PPX and measured PPX-mediated release of P_i_ using a modified molybdate assay ^40^. Whereas polyP that was incubated with α-synuclein monomers was rapidly hydrolyzed and yielded in the expected amount of P_i_ (Fig. 2g, triangles), presence of 40 μM α-synuclein fibrils protected about 130 μM of P_i_-units against hydrolysis (Fig. 2g). We obtained very similar results when we incubated 40 μM α-synuclein monomers or fibrils with 500 μM polyP, spun down polyP-associated fibrils and measured hydrolyzable polyP in both supernatant and pellet. Over 95% of PPX-hydrolyzable polyP was found in the supernatant of samples containing soluble α-synuclein monomers. In contrast, about 45% of the total polyP pelleted with α-synuclein fibrils, of which about two thirds (~130 μM) were resistant towards PPX-mediated hydrolysis (Fig. 2h). We concluded from these results that α-synuclein-fibrils resist conformational rearrangements by interfering with exopolyphosphatase-mediated polyP hydrolysis.

### Extracellular polyP prevents intracellular enrichment of α-synuclein fibrils

Our findings that polyP associates with pre-formed α-synuclein fibrils and changes their conformation, served to explain results of our previous studies, which showed that α-synuclein fibrils lose their cytotoxicity as soon as polyP is added ^30^. However, they did not explain how polyP is able to protect against amyloid toxicity. We reasoned that polyP might reduce the formation of cytotoxic oligomers by stabilizing the fibrils in a conformation that has previously been shown to be less prone to shedding ^30^. Alternatively, we considered that binding of polyP to the fibrils might either directly or indirectly interfere with the membrane association of α-synuclein ^41, 42^ and/or its cellular uptake ^25, 43^. Lastly, it was also conceivable that polyP binding might increase the turnover of internalized α-synuclein or its sequestration into non-toxic deposits. To gain insights into the potential mechanism(s) by which polyP protects neuronal cells against amyloid toxicity, we compared uptake and intracellular fate of exogenously added α-synuclein fibrils in the absence and presence of polyP. We labeled α-synuclein with AlexaFluor 488, formed mature fibrils, pelleted them by centrifugation and sonicated the fibrils to obtain a mixture of oligomeric species, protofibrils and short mature fibrils (i.e., α-syn^PFF-AF488^) ^44, 45^ (Supplemental Fig. 1a). We confirmed that sonication does not affect the interaction of fibrils with polyP (Supplemental Fig. 1b). We then incubated differentiated SH-SY5Y neuroblastoma cells with α-syn^PFF-AF488^ or freshly prepared fluorescently labeled monomeric α-synuclein (i.e., α-syn^mon-AF488^) at either 4°C or 37°C in the absence or presence of polyP for 24h, and analyzed AF488-fluorescence using confocal microscopy. In the absence of polyP, we detected significant intracellular fluorescence upon incubation of the cells with either α-syn^mon-AF488^ or α-syn^PFF-AF488^ at 37°C but not at 4°C (Fig. 3a). Moreover, we noted an apparently stable association of α-syn^PFF-AF488^ with the cell membrane at both temperatures (Fig. 3a), which was confirmed by trypan blue staining (Supplemental Fig. 2a). These results were fully consistent with previous studies, which reported that both monomers and fibrils use a temperature-sensitive endocytic route for their cellular uptake ^46^, and that α-synuclein fibrils stably associate with cell membranes ^43^. Incubation of the cells in the presence of polyP significantly reduced the intracellular fluorescence signal of α-syn^PFF-AF488^ upon incubation at 37°C as well as the membrane-associated signal upon incubation at either temperature (Fig. 3a). This result was distinctly different from monomeric α-syn^mon-AF488^, whose uptake at 37°C was not affected by polyP. These results strongly suggested that polyP negatively influences the membrane association and/or uptake of α-syn^PFF-AF488^.

**Figure 3.**
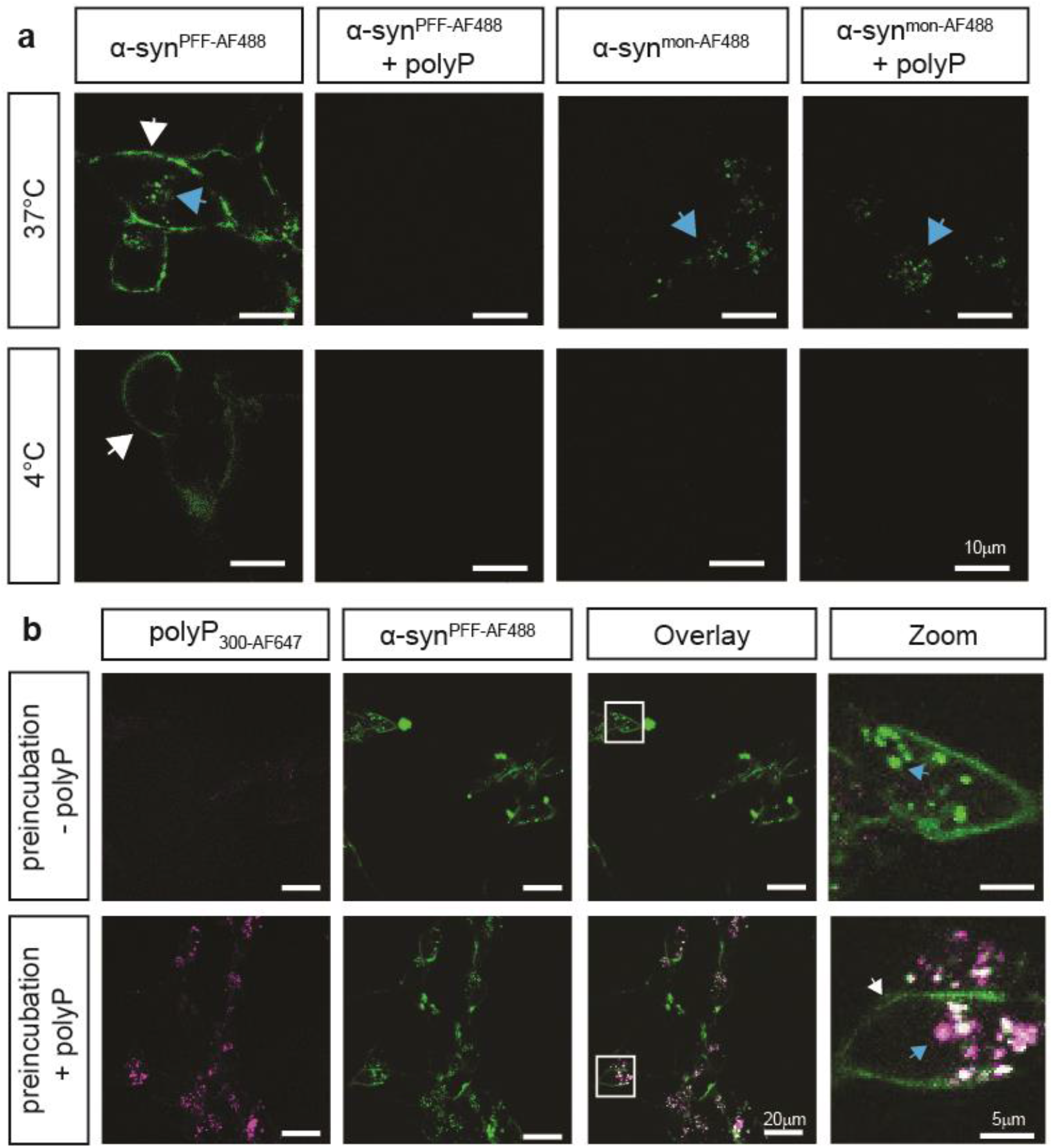
Extracellular polyP prevents intracellular enrichment of α-synuclein fibrils. **a.** Differentiated SH-SY5Y neuroblastoma cells were incubated with 3 μM freshly purified monomeric α-syn^mon-AF488^ or α-syn^PFF-AF488^ fibrils in the absence or presence of 250 μM polyP_300_ (in P_i_-units) at either 4°C or 37°C for 3 h. Membrane-associated α-syn^AF488^ is indicated with white arrows while internalized α-syn^AF488^ is indicated with blue arrows. **b.** Intracellular enrichment of cells with polyP_300_ neither affects uptake nor turnover of α-syn^PFF-AF488^. Differentiated SH-SY5Y cells were incubated with 250 μM polyP_300-AF647_ at 37°C for 24 h and washed to remove extracellular polyP. Then, 3 μM preformed α-syn^PFF-AF488^ fibrils were added and fluorescence microscopy was conducted after 24 h of incubation.

To test whether intracellular polyP influenced the uptake and/or intracellular foci formation of exogenously added α-syn^PFF-AF488^, we incubated differentiated SH-SY5Y neuroblastoma cells with fluorescently-labeled polyP_300_ (i.e., polyP_300_-_AF647_) for 24 h, washed the cells to remove any exogenous polyP and analyzed the cells using a confocal microscope. We observed a clear AF647-fluorescence signal in cells incubated with fluorescently labeled polyP but not in our control cells (Supplemental Fig. 2b). This result confirmed previous studies that showed that neuronal cells are able to take up and enrich exogenous polyP ^37^. When we incubated the polyP-enriched cells with α-syn^PFF-AF488^, we observed the same rapid internalization and intracellular enrichment of α-syn^PFF-AF488^ fibrils that we found in cells that were not pre-treated with polyP (Fig. 3b). We concluded from these experiments that polyP needs to be present in the extracellular space to interfere with the uptake of α-syn^PFF^, and that intracellular polyP does not substantially affect the fate of internalized α-synuclein fibrils. This is despite the fact that we observed a clear co-localization between internalized α-syn^PFF-AF488^ and intracellular polyP_300-AF647_ in select intracellular foci, demonstrating that polyP associates with α-syn^PFF-AF488^ also in the context of intact cells (Fig. 3b, blue arrow).

### PolyP interferes with α-synPFF membrane association

In order to further investigate the influence of polyP on fibril uptake, we incubated differentiated SH-SY5Y cells with α-syn^PFF-AF488^ as before, and either left them untreated or added polyP defined time points after start of the incubation. We reasoned that determining the effects of polyP on cells that contained both membrane-associated and internalized α-syn^PFF –AF488^ would likely reveal at what stage polyP acts. Before the imaging, we washed the cells to remove any unbound α-syn^PFF^ and/or polyP. As expected, incubation of SH-SY5Y neuroblastoma cells with α-syn^PFF-AF488^ in the absence of polyP revealed a persistent association of labeled α-syn^PFF-AF488^ with the cell membrane, and a steady increase in the intracellular fluorescent signal (Fig. 4a). When we added polyP to cells that were pre-incubated with α-syn^PFF-AF488^ for 2 hours, and imaged the samples 30 min later, we observed a significantly reduced signal of membrane-associated α-syn^PFF-AF488^ and lower levels of intracellular α-syn^PFF-AF488^ compared to the control cells. In the presence of polyP, the fluorescence signals did not significantly change over the next hours of incubation and only a slight increase in the intracellular signal of α-syn^PFF-AF488^ was observed after 24h of incubation. Addition of polyP at later time points (i.e., 4 or 6 hours) caused a similar cessation in α-syn^PFF-AF488^ uptake, and a decrease in cell membrane-associated α-syn^PFF-AF488^ signal (Fig. 4a). Upon addition of fluorescently labeled polyP_300-AF647_ to cells pretreated with α-syn^PFF-AF488^ for 6 h, we found both fluorescence signals to co-localize on the outside of the cells, consistent with the formation of polyP-fibril complexes (Fig. 4b). These results strongly suggested that binding of polyP to α-synuclein fibrils interferes with the membrane association of α-synuclein, and hence prevents the uptake of fibrils. They also served to explain why the uptake of monomeric α-synuclein, which does not stably interact with polyP, is unaffected by the presence of polyP.

**Figure 4:**
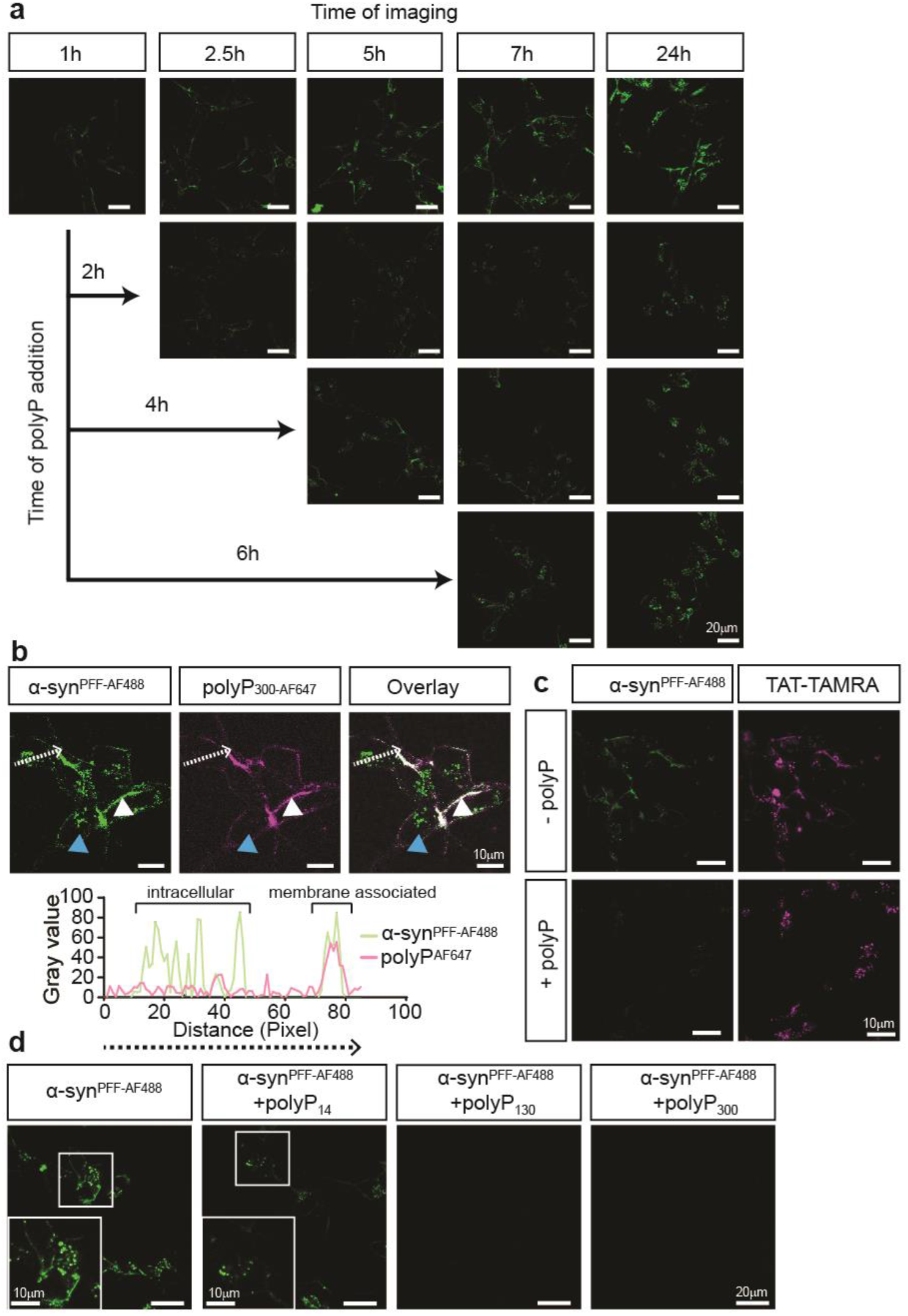
PolyP prevents the association of α-synuclein fibrils with the membrane. **a.** Uptake of 3 >μM preformed α-syn^PFF-AF488^ fibrils after 1, 2.5, 5, 7 and 24 hours into differentiated SH-SY5Y cells. After 2, 4 or 6 hours of incubation, 250 μM polyP_300-AF647_ was added, and the uptake of α-syn^PFF-AF488^ fibrils was monitored as indicated. **b.** Upper panel: Differentiated SH-SY5Y cells were incubated with 3 μM preformed α-syn^PFF-AF488^ fibrils for 6 hours. Then, 250 μM polyp 300-AF647 was added, and co-localization of α-syn^PFF-AF488^ fibrils and polyP_300-AF647_ was determined. The intracellular α-syn^PFF-AF467^ signal is indicated with blue arrows while extracellular α-syn^PFF-AF467^ is indicated with white arrow. polyP_300-AF647_ was only detected on the cell surface. Lower panel: Fluorescence signal of α-syn^PFF-AF488^ fibrils and polyP_300-AF647_ as measured along the white line marked in the upper figure using the plot profile analysis in ImageJ. **c.** Differentiated SH-SY5Y cells were incubated with 5 μM TAT-TAMRA and 3 μM α-syn^PFF-AF488^ in the presence or absence of 250 μM polyP_300_. After 3 hours of incubation, the uptake was monitored. **d.** Differentiated SH-SY5Y cells were incubated with α-synuclein fibrils for 24 hours in the absence or presence of 250 μM of different chain lengths of polyP.

Recent studies suggested that one mechanism by which α-syn^PFF-AF488^ enter cells is through the interaction with heparin glycan receptors ^47^, in a mechanism termed micropinocytosis ^48^. To investigate the possibility that polyP inhibits the uptake of α-syn^PFF-AF488^ by generally blocking micropinocytosis, we monitored the influence of polyP on the uptake of the Trans-Activator of Transcription (TAT) protein fused to the fluorescent dye TAMRA (TAT-TAMRA) (AnaSpec). TAT is a small viral protein, which contains the heparin sulfate binding sequence necessary for its internalization via micropinocytosis ^49, 50^. We incubated differentiated SH-SY5Y cells with both TAT-TAMRA and α-syn^PFF-AF488^ either in the absence or in the presence of polyP_300_ and monitored the uptake of both proteins via fluorescence microscopy. In the absence of polyP_300_, we observed signals for both α-syn^PFF-AF488^ and TAT-TAMRA in the cells, indicating that both proteins were taken up (Fig. 4c). In the presence of polyP, however, we observed only the TAT-TAMRA signal inside the cells (Fig. 4c). These results are consistent with the model that polyp selectively prevents the uptake of fibrillary α-synuclein without generally interfering with endocytosis mechanisms.

To finally test whether the chain lengths of polyP influences its ability to prevent uptake of α-syn^PFF-AF488^, we incubated differentiated SH-SY5Y cells with α-syn^PFF-AF488^ as before but added 250 μM (in P_i-_units) of either polyP_14_, polyP_130_ or polyP_300_. Analysis of internalized α-syn^PFF-AF488^ after 24 hours demonstrated that whereas the longer polyP-chains completely inhibited the uptake of α-syn^PFF-AF488^, presence of polyP_14_ had a much-diminished effect on the uptake (Fig. 4d). These results were in excellent agreement with our previous competition studies that showed that polyP_14_ chains are substantially less effective in binding to α-syn^PFF-AF488^ and/or competing with polyP_300_ and excluded that the observed effects are simply due to the presence of densely charged polyanions. Instead, these results provided supportive evidence for the conclusion that the mechanism by which polyP protects neuronal cells against α-synuclein toxicity is through its specific interactions with extracellular α-synuclein fibrils, effectively preventing their association with the cell membrane and limiting their uptake into neuronal cells.

## DISCUSSION

### Effects of polyP on α-synuclein fibril formation and structure

Previous work from our lab demonstrated that polyP effectively accelerates fibril formation of disease-related amyloids, and protects against amyloid toxicity both in cell culture as well as in disease models of *C. elegans* ^30^. To gain insights into the mechanism by which polyP exerts these effects, we tested at what stage during the fibril-forming process, polyP acts on α-synuclein, one of the major amyloidogenic proteins involved in Parkinson’s Disease. These studies showed that polyP is unable to accelerate the rate-limiting step of α-synuclein fibril formation. Instead, polyP binds to α-synuclein species that begin to accumulate at the end of the lag phase, and are present throughout the elongation and stationary phase of fibril formation. These results agreed well with previous solution studies, which showed that polyP does not promote the conversion of α – helical proteins into β - sheet structures but instead stabilizes folding intermediates once they have adopted a β-sheet conformation ^51^. These results also make physiological sense as they exclude the possibility that simple co-presence of polyP and α-synuclein in the same cellular compartment cause fibril formation.

Earlier work on α-synuclein has shown that the primary nucleation step involves structural changes within α-synuclein monomers and formation of small pre-fibrillar oligomeric intermediates, which are rich in β-sheet structures yet unable to increase ThT-fluorescence ^10, 38, 52^. It has been proposed that these oligomers undergo a cooperative conformational change, leading to the formation of ThT-positive proto-fibrils and fibrils ^52^. Our findings that polyP-binding slightly yet reproducibly precedes ThT binding and substantially accelerates the formation of ThT-positive protofibrils and fibrils suggest that polyP serves as a binding scaffold for pre-fibrillar oligomers and increases the cooperativity of fibril formation.

A recently solved cryo-EM structure of mature α-synuclein_1-121_ fibrils revealed that the double-twisted nature of the fibrils results from the association of two protofilaments, which are stabilized by intermolecular salt bridges ^53^. Moreover, the fibrils are characterized by dense positively charged patches that are located in the vicinity of the interface and run in parallel to the fibril axis. We now hypothesize that binding of the negatively charged polyP chains to such densely positively charged patches that run alongside individual oligomers will support the correct orientation of the oligomers along the fiber axis hence nucleate fibril formation. This model would explain why polyP shows very low apparent affinity for soluble α-synuclein monomers, and provide some rationale for the very low binding stoichiometry of polyP to α-synuclein, which is a mere 5 P_i_-units per one α-synuclein monomer. However, future high-resolution structure studies are clearly necessary to answer the important question as to how polyP and α-synuclein fibrils interact.

Our studies showed that polyP does not only change the morphology of α-synuclein fibrils when present during *de novo* fibril formation but also of mature α-synuclein fibrils. These results agree with recent findings, which suggested that fibrils are intrinsically dynamic and can adopt different conformations over a time-scale of weeks to months ^54^. In the case of polyP, however, α-synuclein fibril conformation dramatically changes within less than 20 min, clearly indicating that changes in fibril morphology happen on a much shorter time scale. Future work needs to be done to investigate the relative effects of polyP on the fibril morphology of disease-associated mutant, which appear to be more resistant to morphology changes than wild-type α-synuclein. In either case, however, our results suggest that polyP might play a pivotal role as modifier of disease-associated fibrils.

### Mechanistic insights into polyP’s protective role against α-synuclein toxicity

The toxicity associated with α-synuclein fibril formation has long been attributed to the cellular accumulation of insoluble fibril deposits ^55, 56^. However, increasing evidence now suggests that oligomeric intermediates, which accumulate during amyloid fibril formation, interfere with membrane integrity, mitochondrial activity and/or other physiologically important functions, and elicit the neuro-inflammatory responses associated with the disease ^14^. Similarly, disease progression has also been proposed to be the responsibility of amyloid oligomers, which appear to be able to spread from cell to cell in a prion-like manner ^20, 22, 23^. Rodents that were subjected to α-syn^PFF^ injections into the *striatum*, for instance, developed neurodegeneration in the *substantia nigra* ^57, 58^. These results suggested that physiologically relevant amyloid modifiers, such as polyP, which are present in the extracellular space in the brain ^37^, and protect against amyloid-induced cytotoxicity in cell culture models, might be involved in the spreading of the disease. We now revealed that binding of polyP to extracellular α-syn^PFF^ decreased the membrane association of α-syn^PFF^ and significantly reduced the internalization of α-synuclein fibrils (Fig. 5), likely explaining the reason for polyP’s cytoprotective effects. We found this effect to be highly specific for amyloid fibrils since neither the uptake of α-synuclein monomers nor of TAT-TAMRA, which, like α-syn^PFF^ is internalized via micropinocytosis ^49, 50^, was affected by the presence of polyP. Moreover, shorter polyP-chain lengths, which are much less effective in interacting with α-syn^PFF^ compared to longer chains, were found to be also much less effective in preventing the uptake of the fibrils. Finally, we found that intracellular enrichment with polyP had no effect on the amount of internalized α-synuclein fibrils, indicating that polyP blocks the uptake and not the intracellular turnover rate of α-syn^PFF^. These results suggested that the direct interaction between polyP and α-syn^PFF^, through alterations in fibril conformations and/or the abundance of negative charges associated with α-syn^PFF^, prevents the interactions of α-syn^PFF^ fibrils with the negatively charged lipids on the cell membrane and leads to its dissociation from the membrane ^47^. Given that the toxicity of α-synuclein amyloids is attributed to their ability to bind, penetrate and damage the membrane ^59, 60^, our finding that a physiologically relevant compound affects this process, suggests that polyP might play an important role in the development and/or progression of this disease. Tools need to be developed to quantify, monitor and manipulate extra- and intracellular polyP levels and test the exciting idea that prevention of the reported age-associated polyP decline in mammalian brains ^61^ might serve to delay the onset and/or extent of this devastating disease.

**Figure 5.**
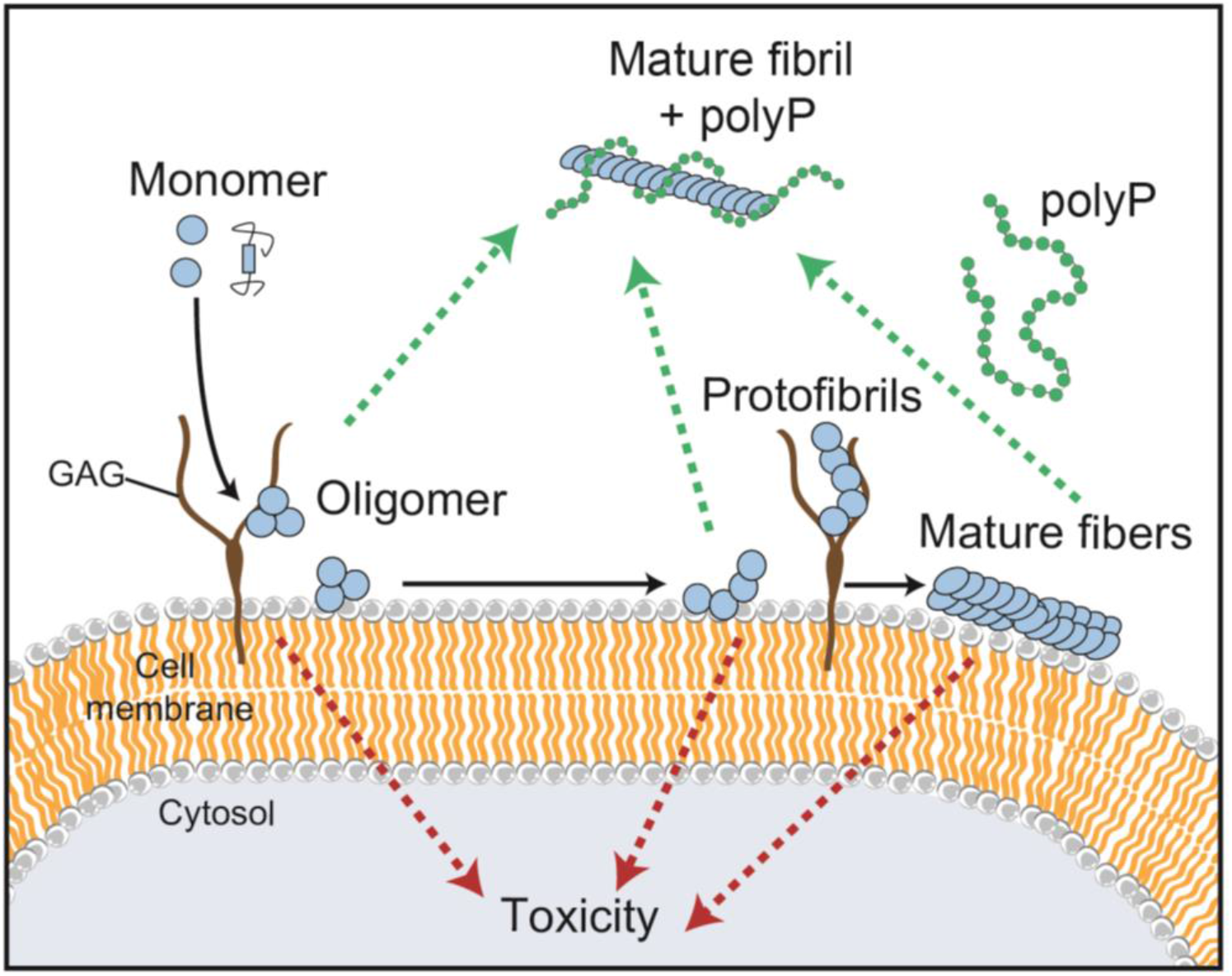
Model for the influence of polyP on fibril formation, morphology and uptake. PolyP accelerates amyloid fibril formation by nucleating pre-fibrillar oligomers. PolyP-associated fibrils have significantly altered fibril morphology. The interaction of polyP with amyloidogenic α-synuclein interferes with their membrane association and there prevents cellular uptake.

## MATERIAL AND METHODS

### PolyP preparation

Defined chain length polyP was a kind gift from Dr. Toshikazu Shiba (Regenetiss, Japan). PolyP was labeled with Alexa Fluor-647 as described ^62^. In brief, 125 μM of polyP_300_ chain was incubated with 2.5 mM Alexa Fluor 647 cadaverine (Life Technologies) and 200 mM 1-ethyl-3-(3-dimethylaminopropyl) carbodiimide (EDAC) (Invitrogen) in water for 1 hour at 60°C. The reaction was stopped on ice and labeled polyP_300-AF647_ was separated from free dye and unlabeled polyP via a NAP-5 column (GE Healthcare) that was equilibrated with 40 mM KPi, pH 7.5. The concentration of polyP was determined via a toluidine blue (TBO) assay ^63^. In this assay, polyP was mixed with 6 mg/l TBO and the absorbance was measured at 530 and 630 nm. The 530 nm / 630 nm absorbance ratio was determined and the concentration was calculated based on a polyP_300_ standard curve.

### Protein purification and labeling of α-synuclein

Alpha-synuclein WT or α-synuclein A90C mutant was purified as described ^30, 64^ with slight modifications. In brief, *E. coli* strain BL21 (DE3) containing the α-synuclein-expressing vector pT7-7 was grown in Luria Broth, supplemented with 200 μg/mL ampicillin until OD_600_ of 0.8-1.0 was reached. The protein expression was induced with 0.8 mM Isopropyl β-D-1-thiogalactopyranoside for 4 h. Then, bacteria were harvested at 4,500 × g for 20 minutes and 4°C. The pellet was resuspended in 50 ml lysis buffer (10 mM Tris-HCl, pH 8.0, 1 mM EDTA, Roche Complete protease inhibitor cocktail) and the lysate was boiled for 15-20 minutes. The aggregated proteins were removed by centrifugation at 13,500 × g for 30 minutes. Next, 136 μl/ml of a 10% w/v solution streptomycin sulfate solution and 228 μl/ml glacial acetic acid were added to the supernatant. After another centrifugation step at 13,500 × g for 30 minutes, the supernatant was removed and mixed in a 1:1 ratio with saturated ammonium sulfate and incubated stirring at 4°C for 1 h. The mixture was spun down at 13,500 × g for 30 minutes and the pellet was resuspended in 10 mM Tris-HCl, pH 7.5. Concentrated NaOH was used to adjust the pH of the suspension to pH 7.5. Afterwards, the protein was dialyzed against 10 mM Tris-HCl pH 7.5, 50 mM NaCl, filtered and loaded onto three connected 5 ml HiTrap Q HP columns (GE Healthcare). Washing steps were performed with 10 mM Tris-HCl pH 7.5, 50 mM NaCl and the protein was eluted with a linear gradient from 50 mM to 500 mM NaCl. The protein-containing fractions were combined and dialyzed against 50 mM ammonium bicarbonate, pH 7.8. Oligomeric α-synuclein species were removed by filtering the protein through a 50-kDa cut-off column (Amicon, Millipore). Aliquots of the protein were prepared, lyophilized and stored at −80°C. For crosslinking of α-synuclein-A90C with Alexa Fluor 488 – maleimide (Invitrogen), 100 μM of the protein was incubated with 1 mM tris(2-carboxyethyl)phosphine (Invitrogen) for 30 min at RT protected from light. Alexa Fluor 488 – maleimide was added in 3-fold excess and the mixture was incubated over night at 4°C. The reaction was stopped by adding 2 mM Dithiothreitol. The free dye was removed using a NAP column (GE Healthcare). The concentration of dye and protein were determined by measuring absorbance at 488 and 280 nm, respectively. Saccharomyces cerevisiae exopolyphosphate (ScPPX) was purified according to Pokhrel et al ^65^ with slight modifications. In brief, MJG317 (BL21 / pScPPX2 = S. cereviseae PPX1 in pET-15b) was incubated overnight at 37°C without shaking in LB containing 100 μg mL-1 ampicillin. The next day the cultures were shaken for about 30 min at 180 rpm at 37°C until they reach an absorbance of 0.4 - 0.5. Additional 100 μg mL-1 ampicillin and isopropyl β-D-1-thiogalactopyranoside (IPTG) to a final concentration of 1 mM were added and protein was expressed by incubating the cells for 4 h at 37 °C with shaking at 180 rpm. The cells were harvested by centrifuging for 20 min at 4000 rpm at 4°C. The pellet was resuspended in 50 mM sodium phosphate buffer, 500 mM NaCl, 10 mM imidazole (pH 8) and 1 mg / mL lysozyme, 2 mM MgCl2, and 50 U / mL Benzonase was added. The solution was incubated for 30 min on ice to digest nucleotides. Cell lyses was performed via sonication with two cycles of 50% power pulsing 5s on and 5 s off for 2 min with 2 min rest between cycles. The protein lysate was centrifuged to remove cell debris for 20 min at 20,000 g at 4°C and loaded onto a nickel-charged chelating column. After washing with 50 mM sodium phosphate buffer, 0.5 M NaCl, 10 mM imidazole (pH 8) and 50 mM sodium phosphate buffer, 0.5 M NaCl, 20 mM imidazole (pH 8) samples were eluted with 50 mM sodium phosphate buffer, 0.5 M NaCl, 0.5 M imidazole (pH 8). Fractions containing ScPPX were pooled and dialyzed twice against 2L of 20 mM Tris-HCl (pH 7.5), 50 mM KCl, 30% (v/v) glycerol. Precipitated protein was removed via centrifugation for 20 min at 20,000 g at 4°C, 50% glycerol was added and the protein was stored at −80°C.

### Preparation of fluorescently labeled α-syn^PFF^

To generate α-syn^PFF-AF488^, 760 μM freshly purified α-synuclein monomers were incubated with 40 μM labeled α-synuclein-AF488 in 40 mM KPi, 50 mM KCl, pH 7.5 for 24 hours at 37°C under continuous shaking using two 2 mm borosilicate glass beads (Aldrich) in clear 96-well polystyrene microplates (Corning) ^66^. Samples from the 96 well plate were combined in Eppendorf tubes and collected via centrifugation at 20,000 × g, 20 min, RT and the pellets were washed twice with 40 mM KPi, 50 mM KCl, pH 7.5 to remove smaller oligomers. After the final spin, the pellets were resuspended in 40 mM KPi, 50 mM KCl, pH 7.5 and sonicated 3 × 5 seconds on ice with an amplitude of 50%. The concentration of fibrils was determined by incubating a small aliquot of α-syn^PFF-AF488^ in 8 M urea, 20 mM Tris pH 7.5, measuring the absorbance at 280 nm and calculating the concentration with the extinction coefficient of 5960 L mol-1 cm-1. Aliquots were taken and stored at – 80°C.

### Thioflavin T fluorescence and fluorescence polarization (FP) measurements

Freshly purified α-synuclein monomers (concentrations provided in the respective figure legends) were incubated with 10 μM thioflavin T (ThT; Sigma) in 40 mM KPi, 50 mM KCl, pH 7.5 at 37°C and two 2 mm borosilicate glass beads (Aldrich) in the absence or presence of polyP_14_ or polyP_300_ (given in P_i_ units). For ThT-measurements, samples were pipetted into black 96-well polystyrene microplates with clear bottoms (Greiners). ThT fluorescence was detected in 10 min intervals using a Synergy HTX MultiMode Microplate Reader (Biotec) using an excitation of 440 nm, emission of 485 nm and a gain of 35. To monitor the binding of polyP to α-synuclein during fibril formation, samples were pipetted into 96-well polystyrene microplates with clear bottoms (Greiners). Fluorescence polarization was measured in a Tecan Infinite M1000 Microplate reader, using an excitation of 635 nm and an emission of 675 nm. Measurements were taken in 10-min intervals.

### PolyP binding and competition assays using anisotropy measurements

Anisotropy measurements were conducted in the Varian Cary eclipse Fluorescence Spectrophotometer, using an excitation of 640 nm and an emission of 675 nm (PMT value set between 50 and 100). Samples containing 50 μM polyP-AF647 in 40 mM KPi, 50 mM KCl, pH 7.5 at 37°C. At the indicated time points, 30 μM of α-synuclein monomers or α-synuclein fibrils were added and anisotropy was further monitored over time. For competition experiments, α-synuclein fibrils were formed in the presence of polyP_300_-_AF647_ as before. At defined time points, unlabeled polyP_14_ or polyP_300_ was added, and the anisotropy signal was monitored over time.

### Negative stain of fibrils and transmission electron microscopy (TEM) analysis

To form fibrils for TEM analysis, 300 μM freshly prepared α-synuclein monomers were incubated either in the absence of polyP (i.e., α-syn^alone^), or in presence of 7.5 mM (per Pi) polyP_300_ (i.e., α-syn^polyP^) for 24 hours at 37°C with 2 mm borosilicate glass beads under continuous shaking. Alpha-syn^alone^ fibrils were then either left untreated or were incubated with 7.5 mM polyP_300_ for 20 min (i.e. α-syn^alone→polyP^). Samples were negatively stained with 0.75% uranyl formate (pH 5.5-6.0) on thin amorphous carbon layered 400-mesh copper grids (Pelco) in a procedure according to ^67^. Briefly, 5 μl of the sample was applied onto the grid and left for 3 min before removing it with Whatman paper. The grid was washed twice with 5 μl ddH_2_O followed by three applications of 5 μL uranyl formate. The liquid was removed using a vacuum. Grids were imaged at room temperature using a Fei Tecnai 12 microscope operating at 120kV. Images were acquired on a US 4000 CCD camera at 66873x resulting in a sampling of 2.21 Å/pixel. About 45 individual α-synuclein filaments were selected across 10 micrographs of each sample and the filament widths were determined using the micrograph dimensions as a reference. Pixel widths were converted into angstroms using the program imageJ.

### X-Ray Fiber Diffraction

Alpha-synuclein fibrils were grown with and without polyP as described above. Prior orientation for diffraction, 1-2 ml of a solution containing 100 μM α-synuclein fibrils were washed 3 times with 10 mM Tris pH 7. Then, fibrils were pelleted by centrifugation (15,000xg, 5min). The supernatant was removed, and the pellet was resuspended in 5-10 μl 10 mM Tris pH 7.0. Then, 5 μl of the fibril pellet was placed between two fire-polished silanized glass capillaries and oriented by air-drying. The glass capillaries with the aligned fibrils were mounted on a brass pin. Diffraction patterns were recorded using 1.13 Å X-rays produced by a 21-ID-D beamline, Argonne Photon Source (APS). All patterns were collected at a distance of 200 mm and analyzed using the Adxv software package ^68^.

### PolyP concentration determination using the molybdate assay

40 μM of α-synuclein monomers or fibrils, prepared in 40 mM Hepes, pH 7.5 and 50 mM KCl, were incubated with the indicated concentrations of polyP_300_ for 10 min at RT in a clear 96-well plate (Corning). The samples were either used directly or spun down at 20,000 × g for 20 minutes at RT to remove any unbound polyP. The pellets were resuspended in 40 mM Hepes (pH 7.5), 50 mM KCl. Next, 8 μg/ml *Saccharomyces cerevisiae* exopolyphosphate (ScPPX) and 1 mM MgCl_2_ was added to each sample and the incubation was continued for 105 min (for spin down) or 120 min (for titration) at RT. To stop the reaction and detect P_i_, 25 μl of a detection solution containing 600 mM H_2_SO_4_, 88 mM ascorbic acid, 0.6 mM potassium antimony tartrate, and 2.4 mM ammonium heptamolybdate was added ^40, 65^. The reactions were developed for 30 min. Then, the precipitated proteins were re-solubilized with 100 μl of 1 M NaOH, and the absorbance was measured at 882 nm using a Tecan M1000 plate reader. The free phosphate concentration was determined with a standard curve of sodium phosphate, which was prepared in parallel with each experiment. After the spun down the phosphate measured in the supernatant was considered free, and the phosphate measured in the pellet was considered bound and unprotected. The bound and protected fraction of phosphate was calculated as total polyphosphate (measured in parallel) minus supernatant phosphate minus pellet phosphate.

### Cell culture experiments and microscopy-

Human neuroblastoma cells SH-SY5Y cells (ATCC CRL-2266) were cultured in Dulbecco’s Modified Eagle Medium: Nutrient Mixture F-12 (Thermo Fisher) medium supplemented with 10% (v/v) heat inactivated fetal bovine serum (Sigma-Aldrich), 1% (w/v) penicillin/streptomycin (Life Technologies) at 37°C and 5% CO_2_. The media was changed every 2–3 days and cells were split 1–2 times per week. For microscopy experiments, 60 000 cells/ml were seeded in 8-well Nunc™ Lab-Tek™ II Chambered Coverglass (Thermo Fisher) and differentiated for 5-7 days by adding 10 μM all-trans retinoic acid (Sigma-Aldrich) every other day. The differentiated cells were incubated with 3 μM α-syn^PFF-AF488^ in the presence or absence of the indicated concentrations of polyP at either 37°C or 4°C for the indicated times. Before the imaging, the media was exchanged to DMEM/F12 without phenol red (Thermo Fisher), supplemented with 10% (v/v) heat inactivated fetal bovine serum (Sigma-Aldrich), 1% (w/v) penicillin/streptomycin (Life Technologies). Cells were imaged using a Leica SP8 high resolution microscope. To distinguish between the inside and outside signals, the cells were treated the same way but 0.05% of the membrane impermeable dye Trypan blue was used for 15 sec prior to the imaging to quench extracellular fluorescence ^43^. To enrich for endogenous polyP, SH-SY5Y cells were seeded and differentiated as described above. Once differentiated, cells were either left untreated or incubated with 250 μM polyP-AF647 (per Pi) for 24 hours. Subsequently, fresh media was added to the cells for 6 hours. Afterwards, cells were incubated with 3 μM α-syn^PFF-AF488^ for 24 hours. As before, the media was changed before imaging and cells were imaged using Leica SP8 high resolution microscope. To test the influence of polyP during the α-syn^PFF-AF488^ uptake, differentiated SH-SY5Y cells were incubated with 3 μM α-syn^PFF-AF488^ at 37°C. After 2, 4 or 6 hours, 250 μM polyP-AF647 was added to the cells. Cells were imaged at time points 1, 2.5, 5, 7 and 24 hours. To test for co-localization of α-syn^PFF-AF488^ and polyP-AF647, cells were incubated with 3 μM α-syn^PFF-AF488^ at 37°C. After 6 h 250 μM polyP-AF647 was added and cells were imaged after 7 h. To monitor the influence of polyP on the uptake of TAT, differentiated SH-SY5Y cells were incubated with 5 μM TAT-TAMRA (AnaSpec) and 3 μM α-syn^PFF-AF488^ either in the presence or in the absence of 250 μM polyP. After 3 hours of incubation, the cells were imaged with a Leica SP8 high resolution microscope.

### Statistics

Two-tailed Students t-tests were performed when two groups were compared. One-way ANOVA was performed when comparing more than two groups. P-values under 0.05 were considered significant. All data in bar charts are displayed as mean +/- SD. Replicate numbers (n) are listed in each figure legend. Prism 7.04 (GraphPad) was used to perform statistical analysis. Annova analysis in figure 2 b shows an F value of 8.435 and a degree of freedom of 3.

## Acknowledgments

Defined length polyP chains were kindly provided by T. Shiba (Regenetiss, Japan) and the Morrissey lab. We thank the ATCC for providing us with SH-SY5Y cells. This work was funded by the NIH grants GM122506 to U.J. and a grant from the American Parkinson Disease Association to M.I.I. and A.S. Ju.L. was funded by a scholarship from the Boehringer Ingelheim Foundation.

## Data availability statement

The datasets generated are available from the corresponding author upon request.

## Author contribution

JL: Designed and performed research, analyzed data, wrote the paper

ET: Performed research, analyzed data

DS: Designed and directed research JAL: Performed research

MII: Designed and performed research, analyzed data

AS: Designed and performed research, analyzed data

NY: Designed and performed research, analyzed data

PH: Performed research

UJ: Designed and directed the project, analyzed data, wrote paper

